# Familial Danish dementia young Knock-in rats expressing humanized APP and human Aβ show impaired pre and postsynaptic glutamatergic transmission

**DOI:** 10.1101/2021.06.24.449787

**Authors:** Tao Yin, Wen Yao, Kelly A. Norris, Luciano D’Adamio

## Abstract

Familial British and Danish dementia (FBD and FDD) are two neurodegenerative disorders caused by mutations in the *Integral membrane protein 2B (ITM2b)*. BRI2, the protein encoded by *ITM2b*, tunes excitatory synaptic transmission at both pre- and post-synaptic terminus. Too, BRI2 interacts with and modulates proteolytic processing of Amyloid-β precursor Protein (APP), whose mutations cause familial forms of Alzheimer disease (FAD). To study pathogenic mechanism triggered by the Danish mutation we generated rats carrying the Danish mutation into the rat *Itm2b* gene (*Itm2b*^*D*^ rats). Given the BRI2/APP interaction and the widely accepted relevance of human Aβ, a proteolytic product of APP, to AD, *Itm2b*^*D*^ rats were engineered to express two humanized *App* alleles, to produce human Aβ. Here, we studied young *Itm2b*^*D*^ rats to investigate early pathogenic changes. We found that peri-adolescent *Itm2b*^*D*^ rats present subtle changes in human Aβ levels along with decreased spontaneous glutamate release and AMPAR-mediated responses but increased short-term synaptic facilitation in the hippocampal Schaeffer-collateral pathway. These changes are like those observed in adult mice producing rodent Aβ and carrying either the Danish or British mutations into the mouse *Itm2b* gene. Collectively, the data show that the pathogenic Danish mutation alters the physiological function of BRI2 at glutamatergic synapses; these functional alterations are detected across species and occur early in life. Future studies will be needed to determine whether this phenomenon represents an early pathogenic event in human dementia.

## INTRODUCTION

Model organisms that reproduce the pathogenesis of human diseases are useful to dissect disease mechanisms, identify therapeutic targets and test therapeutic strategies. Because genetic manipulation has been easier in mice, mice have overtaken rats as the major rodent-based model organism in neurodegeneration research. Thus, to study FDD and FBD, fifteen years ago we generated mice carrying the pathogenic Danish and British dementia mutations (*Itm2b*^*D*^ and *Itm2b*^*B*^ mice) into the *Itm2b* mouse gene (1–3). We choose a knock in (KI) approach rather than the more common transgenic overexpression approach for several reasons. KIs mimic the genetic of FDD and FBD and make no assumption about pathogenic mechanisms (except the unbiased genetic one), while the transgenic approach aims to reproduce pathology (plaques, Neurofibrillary tangles (NFTs), etc.), under the assumption that this “pathology” is pathogenic. In KI models, expression of mutant genes is controlled by endogenous regulatory elements, avoiding issues related to over-expression of disease-proteins in a non-physiological quantitative-spatial-temporal manner. Finally, potential confounding “insertion” effects of transgenes are avoided.

Because rats are better suited to study neurodegenerative diseases, we took advantage of recent developments in geneediting technologies and introduced the familial Danish mutation into the genomic *Itm2b* rat locus (*Itm2b*^*D*^ rats). The rat was the organism of choice for most behavioral, memory and cognitive research -which is critical when studying neurodegenerative diseases-because physiological processes are similar in rats and humans and the rat is an intelligent and quick learner (4–7).

Several procedures that are important in dementia research are more easily performed in rats as compared to mice due to the larger size of the rat brain. Cannulas -to administer drugs, biologics, viruses etc.- and micro-dialysis probes -for sampling extracellular brain levels of neurotransmitters, Aβ, soluble tau etc.- can be accurately directed to individual brain regions, causing less damage and increasing specificity. *In vivo* brain imaging techniques, such as MRI (8) and PET (9–11), can assess the extent and course of neurodegeneration with better spatial resolution in rats. Moreover, rats are large enough for convenient *in vivo* electrophysiological recordings or serial sampling of cerebrospinal fluid for detection of biomarkers.

Finally, gene-expression differences suggest that rats may be advantageous model of neurodegenerative diseases over mice. For example, alternative spicing of *tau* (12–15), which forms NFTs and is mutated in Frontotemporal Dementia (16–23), leads to expression of tau isoforms with three or four microtubule binding domains (3R and 4R, respectively). Adult human and rat brains express both 3R and 4R tau isoforms (24): in contrast, adult mouse brains express only 4R tau(25), suggesting that the rat may be a better model organism for dementias with tauopathy, such as FDD and FBD.

BRI2 physically interacts with and modulates processing of APP, which bears relevance to AD pathogenesis (26–30). In addition, APP processing mediates LTP and memory deficits of Danish and British KI mice (31–36). Aggregated forms of Aβ, a product of APP processing, are by and large considered the main pathogenic molecule in AD. Rat and human APP differ by 3 amino-acids in the Aβ region: given that human Aβ are believed to have higher propensity to form toxic Aβ species as compared to rodent Aβ, we produced rats carrying the humanized Aβ sequence *(App*^*h*^ rats) (37,38). Thus, to study possible interactions between the Danish mutation and human Aβ, *Itm2b*^*D*^ rats were backcrossed to *App*^*h*^ rats. Hence, all rats used in this study produce human and not rodent Aβ species.

Here, we studied peri-adolescent *Itm2b*^*D*^ rats, with the purpose of investigating early dysfunctions that may underlie initial pathogenic mechanisms leading to dementia later in life.

## RESULTS

### Generation of *Itm2b*^*D*^ KI rats carrying humanized *App*^*h*^ alleles

The knock-in founder F0-*Itm2b*^*D*^ rat, which is carrying FDD mutation on *Itm2b* rat gene, was generated by CRISPR/Cas-mediated genome engineering as described in Experimental Procedures and Supporting Information. The F0-*Itm2b*^*D*^ rat, which is a chimera for the *Itm2b* gene, was crossed to WT *(Itm2b*^*w/w*^) Long-Evans rats to generate F1-*Itm2b*^*D/w*^ rats. F1-*Itm2b*^*D/w*^ rats were crossed to WT Long-Evans to generate F2-*Itm2b*^*D/w*^ rats. These crossing were repeated three more times to obtain F5-*Itm2b*^*D/w*^ rats. The probability that F5 rats carry unidentified off-target mutations (except those, if present, on Chr. 15) is ~1.5625%. Male and female F5-*Itm2b*^*D/w*^ rats were crossed to obtain *Itm2b*^*D/w*^, *Itm2b*^*D/D*^ and *Itm2b*^*w/w*^ rats.

The FDD mutation consist of a 10 nucleotides duplication one codon before the normal stop codon (39). This produces a frameshift in the BRI2 sequence generating a precursor protein 11 amino acids larger-than-normal (Figure 1A). To verify that the Danish mutation was correctly inserted into *Itm2b* exon 6, we amplified by PCR the *Itm2b* gene exon 6 *Itm2b*^*D/w*^, *Itm2b*^*D/D*^ and *Itm2b*^*w/w*^ rats. Sequencing of the PCR products shows that the Danish mutation was correctly inserted in the *Itm2b* gene exon 6 (Figure 1B) and encoded for the COOH-terminus of the Danish BRI2 mutant. When we generated FDD KI mice, we humanized the mouse COOH-terminal region of BRI2 by introducing an alanine (A) was substituted for threonine (T) at codon 250 (3). Since that humanization did not result into deposition of ADan peptides in amyloid plaques in KI mice (3), that modification was not repeated in rats.

**Figure 1.**
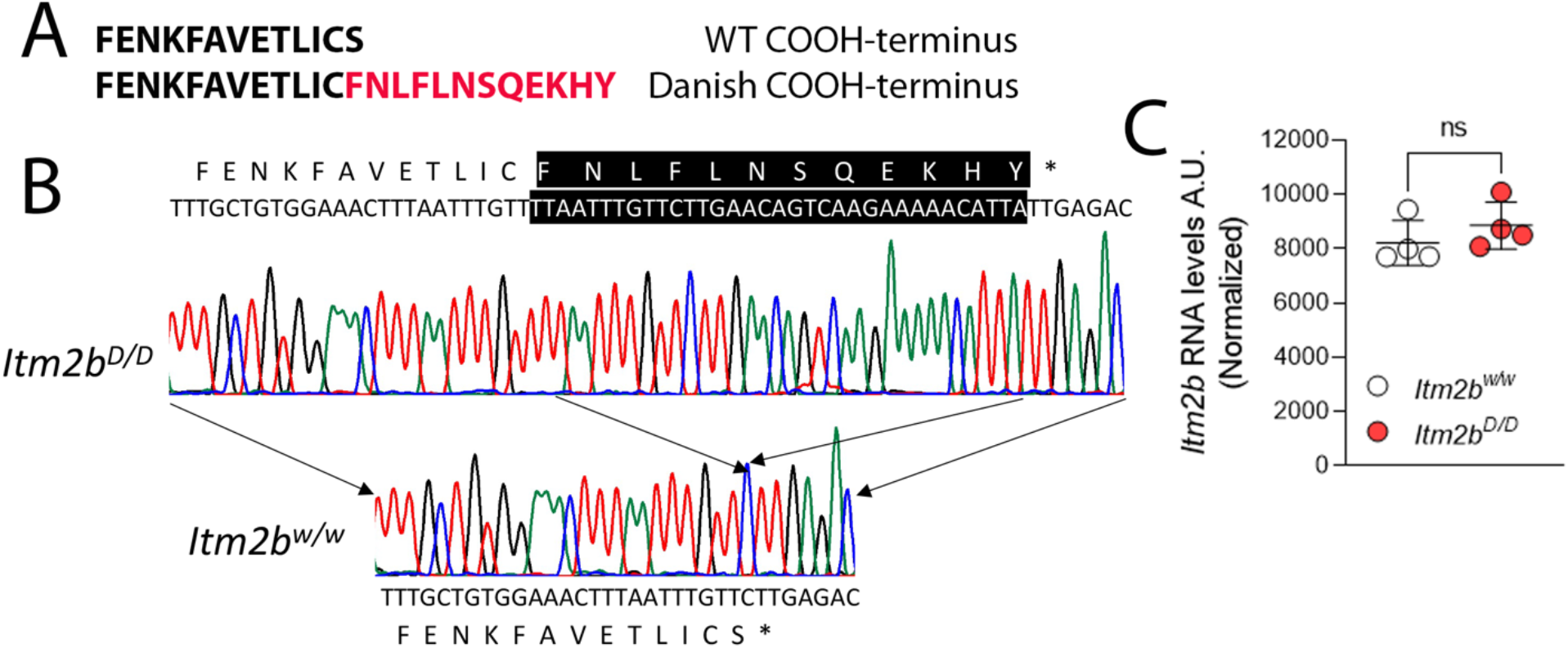
Characterization of *Itm2b*^*D*^ KI rats. (A) Sequences of the COOH-terminus of Bri2-23 (WT) and Bri2-ADan (Danish). (B) PCR amplification and sequencing of the *Itm2b* gene exon 6 from *Itm2b*^*w/w*^ and *Itm2b*^*D,D*^ rats shows that the Danish mutation was correctly inserted in the *Itm2b* exon 6 of *Itm2b*^*D,D*^ rats. This mutation causes the predicted frameshift in the BRI2 sequence generating a precursor protein 11 amino acids larger-than-normal coding for the Bri2-ADan mutant protein (partial DNA sequences of WT and Danish exon 6 are shown. Inserted nucleotides are highlighted in black, and the amino-acid sequences are indicated above -for the Danish mutant allele- and below -for the WT allele-the DNA sequences). (C) Levels of *Itm2b* mRNA in brains of 21 days old *Itm2b*^*w/w*^ and *Itm2b*^*D/D*^ rats were determined by Standard-RNAseq analysis. No significant differences between *Itm2b*^*w/w*^ and *Itm2b*^*D/D*^ rats were evident. Data are represented as mean ± SD. Data were analyzed by Student’s t-test. N=4 rats per genotype.

To generate *Itm2b*^*D/w*^, *Itm2b*^*D/D*^ and *Itm2b*^*w/w*^ rats on a background in which rat *App* has a humanized Aβ region, *Itm2b*^*D/w*^ and *App*^*h/h*^ rats were crossed to generate *Itm2kP*^*D/w*^*;App*^*h/w*^ rats. The *App*^*w*^ allele was removed in subsequent crosses. Henceforth, *Itm2b*^*D/D*^, *Itm2b*^*D/w*^ and *Itm2b*^*w/w*^ rats used in this study have an *App*^*h/h*^ background and produce human and not rodent Aβ species.

To determine whether *Itm2b* expression is disrupted by the introduced mutations, we examined *Itm2b* mRNA levels in p21 *Itm2b*^*D/D*^ and *Itm2b*^*w/w*^ rats by standard RNA-Seq analysis on total brain RNA. The mRNA expression of *Itm2b* shows no significant difference between *Itm2b*^*D/D*^ and *Itm2b*^*w/w*^ rats (Figure1C).

### The *Itm2b*^*D*^ allele encodes for a longer Bri2 precursor protein (Bri2-ADan) that accumulates in primary neurons

BRI2 is a type II membrane protein that is synthesized as an immature precursor (imBRI2). imBRI2 is cleaved at the COOH-terminus by proprotein convertase to produce the NH_2_-terminal mature BRI2 (mBRI2) and the 23 amino acid-long COOH-terminal peptide called Bri23 (40). As noted above, in the Danish patients, a frameshift caused by a 10 nucleotides duplication 5’ to the stop codon leads to the synthesis of a BRI2 precursor protein 11 amino acids larger-than-normal (39). Convertase-mediated cleavage of immature Danish BRI2 generates a WT-like mBRI2 and a 34 amino acid long peptide called ADan, which co-deposits with Aβ species in amyloid fibrils in patients. For clarity, we will refer to the wild type imBri2 as Bri2-Bri23, and to the Danish mutant imBri2 as Bri2-ADan.

To determine whether the *Itm2b*^*D*^ allele codes for Bri2-ADan we examined Bri2 expression in total neuronal lysates isolated from male and female 2 months old *Itm2b*^*D/w*^, *Itm2b*^*D/D*^ and *Itm2b*^*w/w*^ rats. However, the Bri2 antibody tested identified many non-specific bands (Figure S1), making a rigorous assessment of Bri2 expression in rat brains challenging.

Analysis of mouse *Itm2b*^*w/w*^ and *Itm2b*^*D/D*^ primary neurons showed that the mBri2/Bri2-Bri23 ratio in *Itm2b*^*w/w*^ primary neurons was significantly higher than the mBri2/Bri2-ADan ratio in *Itm2b*^*D/D*^ primary neurons (41). In addition, lysosomal inhibition caused accumulation of mBri2 but not Bri2-Bri23 in *Itm2b*^*w/w*^ primary neurons; in contrast, both mBri2 and Bri2-ADan accumulated in *Itm2b*^*D/D*^ primary neurons (41). These observations indicated that the Danish mutation reduced maturation of the mutant precursor Bri2 in mouse neurons. Based on these observations, we probed whether primary neurons could be used to assess mBri2, Bri2-Bri23 and Bri2-ADan expression in KI rats. Primary neurons are a simpler system compared to total brain; this, *per se’*, may reduce the number of non-specific bands identified by anti-Bri2 antibodies. Moreover, inhibition of lysosome-mediated degradation of Bri2 species in primary neurons may help identify specific Bri2 molecules. Thus, primary neurons derived from *Itm2b*^*w/w*^ and *Itm2b*^*D/D*^ rat were treated with the lysosomal inhibitor chloroquine and analyzed by Western blot. The anti-Bri2 antibody identified a band of ~34 kDa in all samples, which was increased by chloroquine (Figure 2A and 2C). These observations are consistent with the ~34 kDa corresponding to mBri2. A second band of ~36 kDa was detected in *Itm2b*^*w/w*^ primary neurons (Figure 2A). In contrast, a slightly larger second band (~37 kDa) that was increased by chloroquine treatment, was detected in *Itm2b*^*D/D*^ primary neurons (Figure 2A and 2C). These observations are consistent with the ~36 kDa and ~37 kDa bands corresponding to Bri2-Bri23 and Bri2-ADan, respectively.

**Figure 2.**
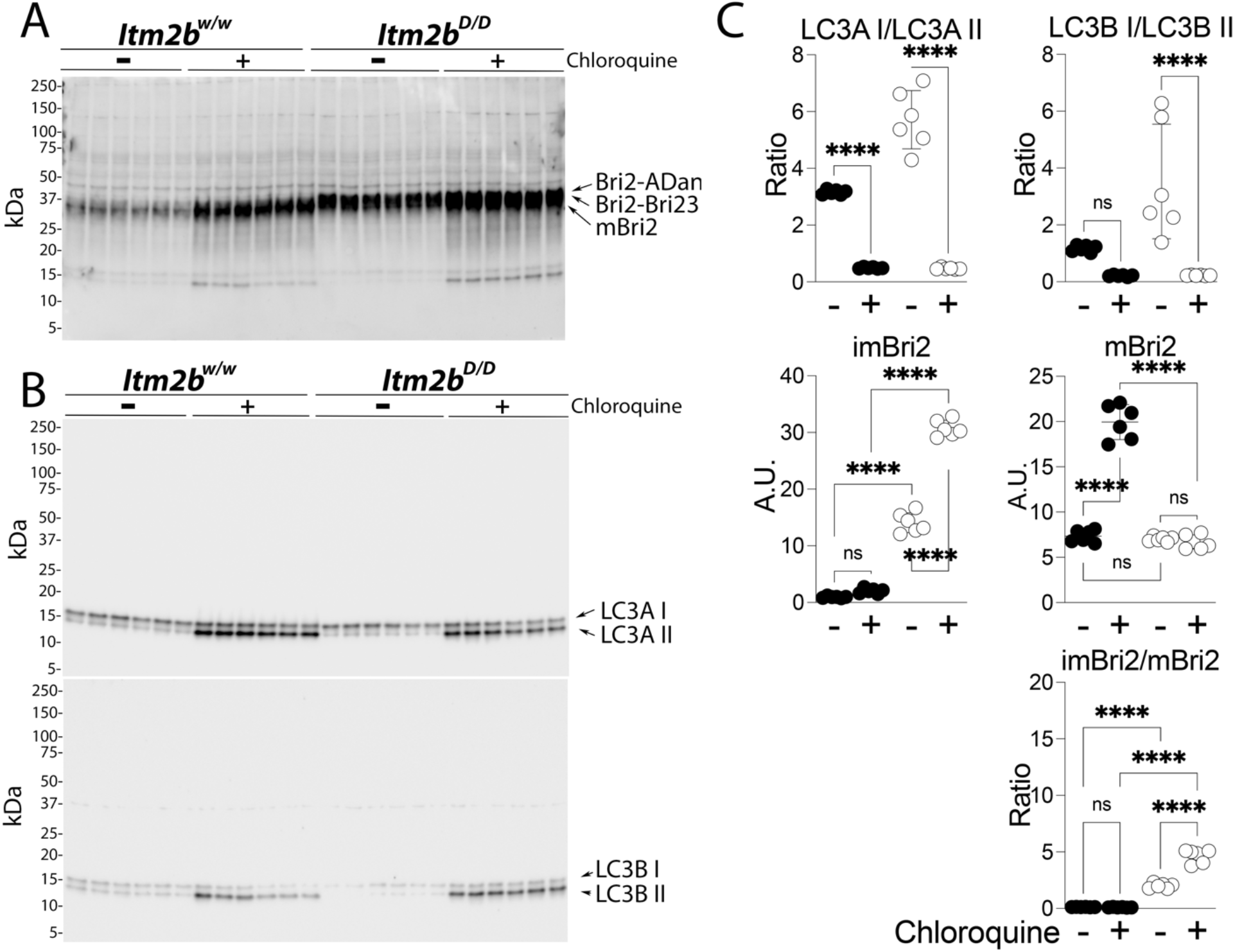
Distinct degradation pathways of Bri2-23 and Bri2-ADan in *Itm2b*^*w/w*^ and *Itm2b*^*D/D*^ primary neurons. WB analysis of Bri2 (A) and LC3A/B (B) from primary hippocampal neurons isolated from *Itm2b*^*w/w*^ and *Itm2b*^*D,D*^ P1 pups treated with 50μM chloroquine for 18h. (C) Quantification of LC3A/B and Bri2 levels. Data are represented as mean ± SD and analyzed by ordinary two-way ANOVA followed by post-hoc Sidak’s multiple comparisons test when ANOVA showed significant differences. ANOVA summary: LC3A: F_interaction (1,20)_=37.36, P<0.0001, F_treatment (1,20)_= 353.5, P<0.0001, F_genotype (1,20)_=36.16, P<0.0001. Post-hoc Sidak’s multiple comparisons test: *Itm2b*^*w/w*^: vehicle (veh) *vs* Chloroquine (Chlor), P<0.0001****, *Itm2b*^*D/D*^: veh *vs* Chlor, P<0.0001****. LC3B: F_interaction (1,20)_=8.11, P<0.01, F_treatment(1,20)_=26.75, P<0.0001, F_genotype (1,20)_=8.283, P<0.01, post-hoc Sidak’s multiple comparisons test: *Itm2b*^*w/w*^: veh *vs* Chlor, P=0.2185, *Itm2b*^*D/D*^: veh *vs* Chlor, P<0.0001****. ImBri2: F_interaction (1,20)_=272.2, P<0.0001, F_treatment (1,20)_= 354.3, P<0.0001 ****, F_genotype (1,20)_= 1966, P<0.0001****, post-hoc Sidak’s multiple comparisons test: *Itm2b*^*w/w*^: veh *vs* Chlor, P=0.5229, *Itm2b*^*D/D*^: veh *vs* Chlor, P<0.0001****. No treatment: *Itm2b*^*w/w*^ *vs Itm2b*^*D/D*^, P<0.0001****. Chlor treatment: *Itm2b*^*w/w*^ *vs Itm2b*^*D/D*^, P<0.0001****. mBri2: F_interaction (1,20)_=207.4, P<0.0001, F_treatment (1,20)_= 186.9, P<0.0001, F_genotype (1,20)_=228.5, P<0.0001, post-hoc Sidak’s multiple comparisons test: *Itm2b*^*w/w*^: veh *vs* Chlor, P<0.0001****, *Itm2b*^*D/D*^: veh *vs* Chlor, P̽0.9966. No treatment: *Itm2b*^*w/w*^ vs *Itm2b*^*D/D*^, P=0.9969 ns. Chlor treatment: *Itm2b*^*w/w*^ *vs Itm2b*^*D/D*^, P<0.0001****. imBri2/mBri2 Ratio: F_interaction (1,20)_=108.3, P<0.0001, F_treatment (1,20)_=104.2, P<0.0001, F_genotype(1,20)_=627.1, P<0.0001, post-hoc Sidak’s multiple comparisons test: *Itm2b*^*w/w*^: veh *vs* Chlor, P>0.9999, *Itm2b*^*D/D*^: veh *vs* Chlor, , P<0.0001****. No treatment: *Itm2b*^*w/w*^ *vs Itm2b*^*D/D*^, P<0.0001****, Chlor treatment: *Itm2b*^*w/w*^ *vs Itm2b*^*D/D*^, P<0.0001****.

Without treatment, the levels of Bri2-ADan in *Itm2b*^*D/D*^ primary neurons were significantly higher than the levels of Bri2-Bri23 in *Itm2b*^*Ww*^ primary neurons (Figure 2A and 2C) and the mBri2/Bri2-Bri23 ratio in *Itm2b*^*w/w*^ primary neurons was significantly higher than the mBri2/Bri2-ADan ratio in *Itm2b*^*D/D*^ primary neurons (Figure 2A and 2C). Chloroquine significantly reduced the LC3A I/LC3A II and LC3B I/LC3B II ratios (Figure 2B and 2C), confirming inhibition of lysosome-mediated degradation.

### Subtle increase in Aβ42 levels in young *Itm2b^D^* KI rats

Sequential processing of APP by α−/γ− secretase and β−/γ−secretase generate the following APP metabolites: sAPPβ, sAPPα, β-CTF, α-CTF, AID/AICD, P3 and Aβ. Since BRI2 interacts with APP and modulates APP processing by α-, β- and γ-secretase (26–30), we determined the steady-state levels of several of these APP metabolites in the central nervous system (CNS) of young male and female *Itm2b*^*D*^ KI rats. Full length APP, α-CTF and β-CTF were measured by Western blot: soluble APPs (sAPPα/sAPPβ) were detected by ELISA, and human Aβ species (Aβ38, Aβ40, Aβ42 and Aβ43) were detected by human Aβ specific-ELISA. These measurements have previously been used for other KI rats generated in our lAb (38,42–44).

Levels of full-length APP, CTFs, Aβ38, Aβ40, Aβ43, sAPPα and sAPPβ were unchanged in 8 weeks old *Itm2b*^*D/w*^, *Itm2b*^*D,D*^ and *Itm2b*^*wlw*^ rats (Figure 3A-C), nor was the Aβ43/Aβ42 ratio altered (Figure 3C). In contrast, there was a slight but significant increase in Aβ42 as well as the Aβ42/Aβ40 ratio in *Itm2b*^*D/D*^ compared to *Itm2b*^*wlw*^ rats (Figure 3C). Small but statistically significant decreases in both Aβ43 and Aβ43/Aβ42 ratio were evident in *Itm2b*^*D/D*^ as compared to *Itm2b*^*w/w*^ rats (Figure 3C). Overall, these data indicate a gene dosage-dependent minor increase in steady-state levels of Aβ42, and decrease in Aβ43, in peri-adolescent *Itm2b*^*D*^ rats. Analysis of older rats will be needed to determine whether the Danish mutation in *Itm2b* alters APP processing in KI rats and whether these alterations may more robustly change the steady-state levels of APP metabolites with aging.

**Figure 3.**
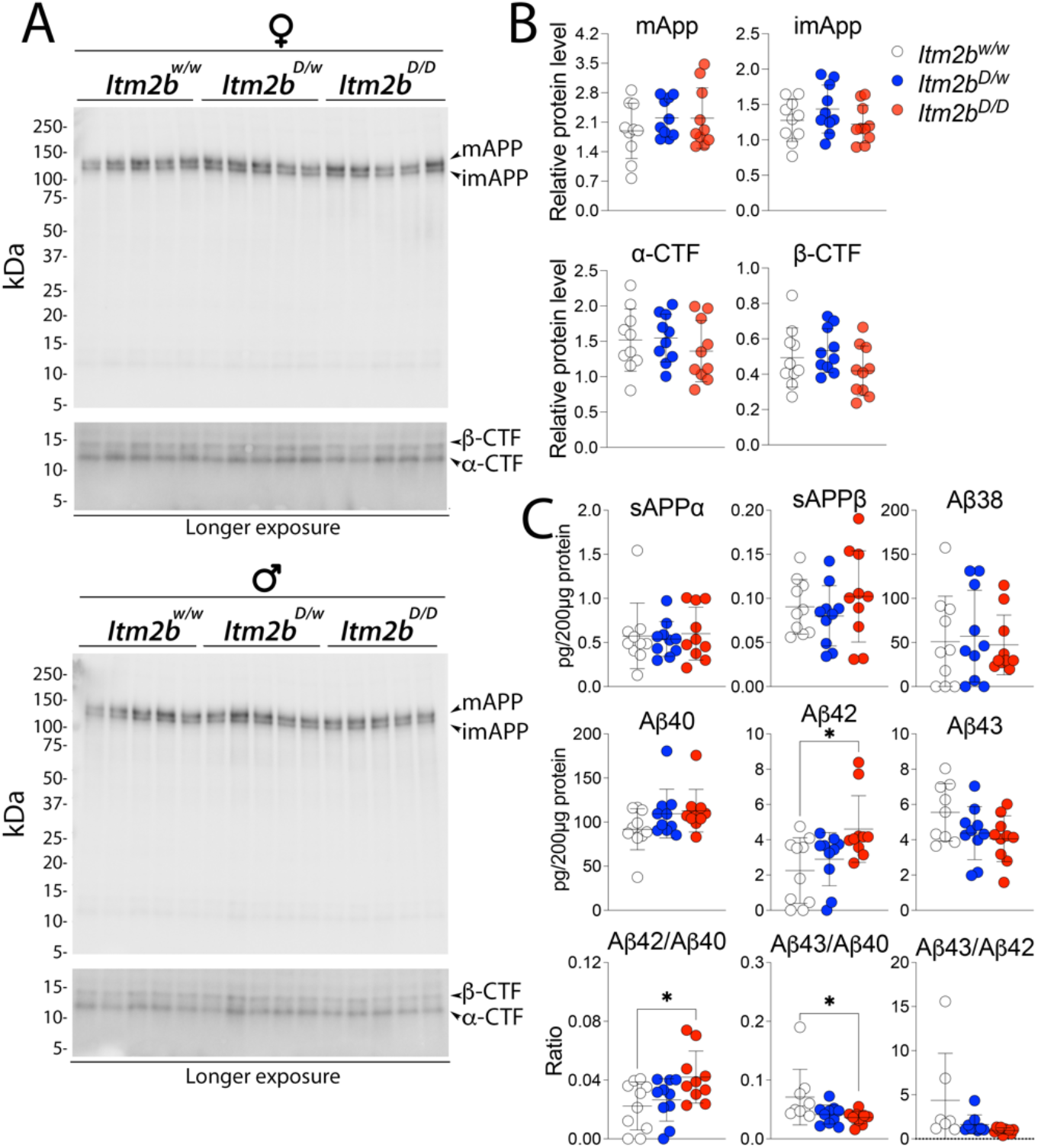
APP metabolite levels in *Itm2b*^*D*^ KI rats. Data are represented as mean ± SD and were analyzed by ordinary one-way ANOVA followed by post-hoc Tukey’s multiple comparisons test when ANOVA showed statistically significant differences. We analyzed 8 weeks old rats, and 5 female and 5 male rats per genotype. (A) Levels of full-length APP, αCTF, and βCTF, were determined by Western analysis of brain lysate of *Itm2b*^*D/D*^, *Itm2b*^*D/w*^ and *Itm2b*^*w/w*^ male and female rats. (B) Quantitation of Western blots. Signal intensity of APP metabolites were normalized to red ponceau staining of nitrocellulose membranes. ANOVA summary: mAPP, F_(2, 27)_ = 0.7931, P=0.4627; imAPP, F_(2, 27)_ = 1.367, P=0.2720; α-CTF, F_(2, 27)_ = 0.6075, P=0.5520; β-CTF, F_(2, 27)_ = 1.614, P=0.2177). (C) Levels of sAPPα, sAPPβ, Aβ38, Aβ40, Aβ42 and Aβ43 were determined by ELISA of brain lysate from the same *Itm2b*^*D/D*^, *Itm2b*^*D/w*^ and *Itm2b*^*w/w*^ male and female rats. [ANOVA summary: sAPPα, F_(2, 27)_ = 0.1084, P=0.8977; sAPPβ, F_(2, 27)_ = 0.7666, P=0.4744; Aβ38, F_(2, 27)_ = 0.1121, P=0.8943; Aβ40, F_(2, 27)_ = 2.030, P=0.1509; Aβ42, F_(2, 27)_ = 4.764, P=0.0169 (post-hoc Tukey’s multiple comparisons test: *Itm2b*^*w/w*^ *vs Itm2b*^*D/w*^, P=0.6966, *Itm2b*^*w/w*^ *vs Itm2b*^*D/D*^, P=0.0159*; *Itm2b*^*D/w*^ *vs Itm2b*^*D/D*^, P=0.0948); Aβ343, F_(2, 26)_ = 2.654, P=0.0893; Aβ42/Aβ40, F_(2,27)_ = 4.074, P=0.0284 (post-hoc Tukey’s multiple comparisons test: *Itm2b*^*w/w*^ *vs Itm2b*^*D/w*^, P=0.8326, *Itm2b*^*w/w*^ *vs Itm2bP*^*D/D*^, P=0.0301*; *Itm2b*^*D/w*^ *vs Itm2b*^*D/D*^, P=0.1022); Aβ43/Aβ40, F_(2, 26)_ = 4.031, P=0.0299 (post-hoc Tukey’s multiple comparisons test: *Itm2b*^*w/w*^ *vs Itm2b*^*D/w*^, P=0.0802, *Itm2b*^*w/w*^*vs Itm2b*^*D/D*^, P=0.0347*; *Itm2b*^*D/w*^ *vs Itm2b*^*D/D*^, P=0.9137); Aβ43/Aβ42, F_(2, 26)_ = 3.281, P=0.0558]. P<0.05*; P<0.01**; P<0.001***; P<0.0001****.

It has been postulated that toxic forms of Aβ are oligomeric (45). Thus, we tested whether toxic oligomers are augmented in peri-adolescent *Itm2b*^*D*^ rats. To this end, we used the prefibrillar oligomer-specific antibody A11 to perform dot blots (46). We found no evidence supporting an increase in neurotoxic brain oligomer levels in peri-adolescent *Itm2b*^*D/w*^ and *Itm2b*^*D/D*^ rats as compared with *Itm2b*^*w/w*^ rats (Figure 4). However, Aβ oligomers appeared to be significantly increased in *Itm2b*^*D/w*^ rats compared to *Itm2b*^*D/D*^ animals (Figure 4). Analysis of older rats will be needed to clarify the relevance of this odd observation.

**Figure 4.**
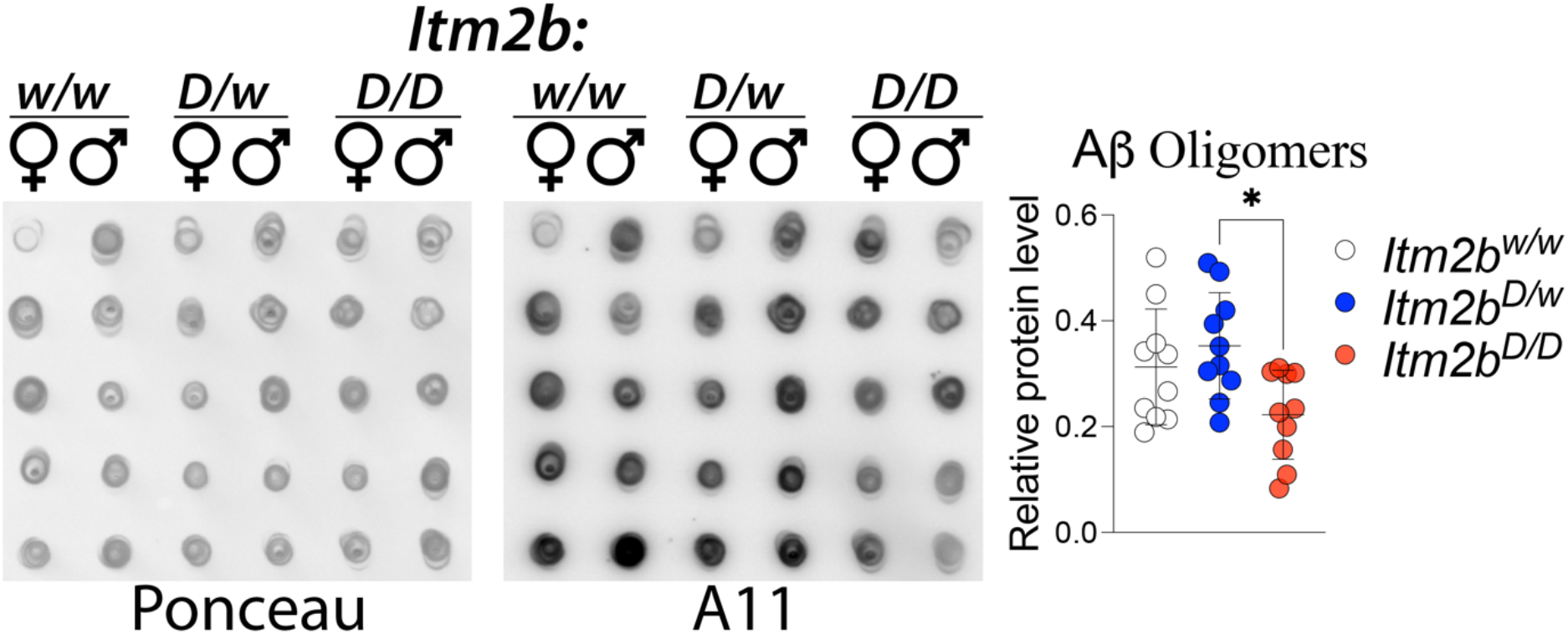
Levels of human Aβ oligomeric species in the brain of peri-adolescent *Itm2b*^*D/D*^, *Itm2b*^*D/w*^ and *Itm2b*^*w/w*^ male and female rats. (A) We analyzed material from the same rats analyzed in Figure 3. Quantitation of dot-blots using the oligomer-specific antibody A11. Before immunoblot analysis, membranes were stained with Ponceau red. Quantitative analysis of A11 blot was normalized to the Ponceau red quantitative analysis. Data are represented as mean ± SD and were analyzed by ordinary oneway ANOVA followed by post-hoc Tukey’s multiple comparisons test when ANOVA showed statistically significant differences. ANOVA summary: F_(2, 27)_ = 4.593, P=0.0192*; post-hoc Tukey’s multiple comparisons test: *Itm2b*^*w/w*^ *vs Itm2b*^*D/w*^, P=0.6406, *Itm2b*^*w/w*^ *vs Itm2b*^*D/D*^, P=0.1195; *Itm2b*^*D/w*^ *vs Itm2b*^*D/D*^, P=0.0170*

### Glutamatergic synaptic transmission at hippocampal SC-CA3>CA1 synapses is impaired in peri-adolescent *Itm2b*^*D*^ rats

Bri2 modulates glutamatergic synaptic transmission at both pre- and post-synaptic termini of Schaeffer-collateral pathway (SC)-CA3>CA1 synapses (47). This function is compromised in both adult *Itm2b*^*D*^ and *Itm2b*^*B*^ KI mice (41). Here, analyzed glutamatergic transmission at SC-CA3>CA1 synapses in young peri-adolescent *Itm2b*^*D*^ male and female rats. First, we analyzed miniature excitatory postsynaptic currents (mEPSC), the frequency of which is determined, in part, by the probability of release (P*r*) of glutamatergic synaptic vesicles release (48). Thus, mEPSC frequency is regulated mostly by pre-synaptic mechanisms: As shown in Figure 5A, B,C, the Danish *Itm2b* mutation caused a significant reduction in the frequency of mEPSC: this reduction is gene-dosage dependent (*Itm2b*^*w/w*^ vs. *Itm2b*^*D,w*^, P=0.003; *Itm2b*^*w/w*^ vs. *Itm2b*^*D/D*^, P<0.0001; *Itm2b*^*D/w*^ vs. *Itm2b*^*D/D*^, P=0.0051) and suggests a decrease in P*r* of glutamatergic synaptic vesicles.

**Figure 5.**
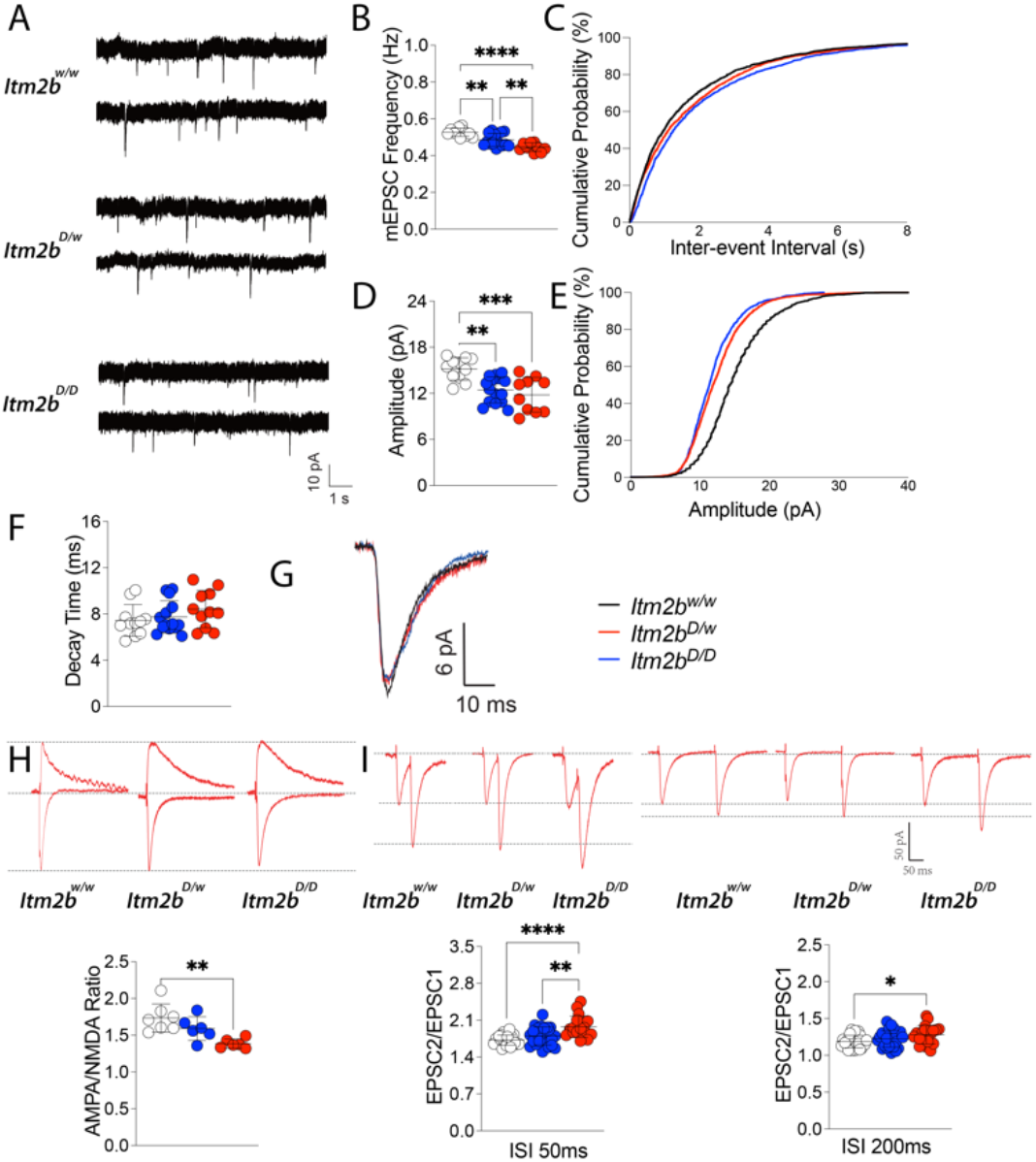
Glutamatergic synaptic transmission is reduced at hippocampal SC-CA3>CA1 synapses of *Itm2b*^*D*^ KI rats. Data are represented as mean ± SD and were analyzed by ordinary one-way ANOVA followed by post-hoc Tukey’s multiple comparisons test when ANOVA showed significant differences. We uswd thw following animals: *Itm2b*^*w/w*^ N=11 (4M/6, 3F/5, indicating that 6 recordings were obtained from the 4 males and 5 recordings from the 3 females), *Itm2b*^*D/w*^ N=15 (3M/7,4F/8), *Itm2b*^*D/D*^ N=10(3M/5,3F/5). (A) Representative recording traces of mEPSC at SC-CA3>CA1 synapses. (B) The Danish mutation causes a significant decrease in mEPSC frequency [ANOVA summary, F_(2, 33)_ = 20.66, P<0.0001****; post-hoc Tukey’s multiple comparisons test: *Itm2b*^*w/w*^ *vs Itm2b*^*D/w*^, P=0.003**, *Itm2b*^*w/w*^ *vs Itm2b*^*D/D*^, P<0.0001****; *Itm2b*^*D/w*^ *vs Itm2b*^*D/D*^, P=0.0051**]. (C) Cumulative probability of AMPAR-mediated mEPSC frequency inter event intervals. (D) [ANOVA summary, F_(2, 33)_ = 10.78, P=0.0002***; post-hoc Tukey’s multiple comparisons test: *Itm2b*^*w/w*^ *vs Itm2b*^*D/w*^, P=0.0016**; *Itm2b*^*w/w*^ *vs Itm2b*^*D/D*^, P=0.0005***; *Itm2b*^*D/w*^ *vs Itm2b*^*D/D*^, P=0.6691]. (E) Cumulative probability of AMPAR-mediated mEPSC amplitude. (F) In contrast, decay time of mEPSC was not significantly changed [ANOVA summary, F_(2, 33)_ = 1.292, P=0.2882]. (G) Average mEPSC of *Itm2b*^*D/D*^, *Itm2b*^*D/w*^ and *Itm2b*^*w/w*^ rats. (H) AMPA/NMDA ratio is significantly decreased in *Itm2b*^*D/D*^ rats [ANOVA summary, F_(2, 16)_ = 8.417, P=0.0032**; post-hoc Tukey’s multiple comparisons test: *Itm2b*^*w/w*^ *vs Itm2b*^*D/w*^, P=0.2393; *Itm2b*^*w/w*^ *vs Itm2b*^*D/D*^, P=0.0023**; *Itm2b*^*D/w*^ *vs Itm2b*^*D/D*^, P=0.0818]. Representative traces are shown on top of the graph (traces are averaged from 20 sweeps). Animals used: *Itm2b*^*w/w*^ N=7 (4M/4,3F/3), *Itm2b*^*D/w*^ N=6 (3M/3,3F/3), *Itm2b*^*D/D*^ N=6 (3M/3,3F/3). (I) Average PPF at 50ms (left panel) and 200ms (right panel) Inter stimulus Interval (ISI) shows that PPF is increased in *Itm2b*^*D/D*^ rats ISI [50ms ISI PPF ANOVA summary: F_(2, 60)_ = 11.89, P<0.0001****; post-hoc Tukey’s multiple comparisons test: *Itm2b*^*w/w*^ *vs Itm2b*^*D/w*^, P=0.2315; *Itm2b*^*w/w*^ *vs Itm2b*^*D/D*^, P<0.0001****; *Itm2b*^*D/w*^ *vs Itm2b*^*D/D*^, P=0.0031**. 200ms ISI PPF ANOVA summary: F_(2, 64)_ = 3.802, P=0.0275*; post-hoc Tukey’s multiple comparisons test: *Itm2b*^*w/w*^ *vs Itm2b*^*D/w*^, P=0.4504; *Itm2b*^*w/w*^ *vs Itm2b*^*D/D*^, P=0.0205*; *Itm2b*^*D/w*^ *vs Itm2b*^*D/D*^, P=0.2462]. Representative traces are shown on top of the panels. P<0.05*; P<0.01**; P<0.001***; P<0.0001****. Animals used: *Itm2b*^*w/w*^ N=18 (4M/10,3F/8), *Itm2b*^*D/w*^ N=20 (3M/10,4F/10), *Itm2b*^*D/D*^ N=17 (3M/8,3F/9).

The amplitude of mEPSC is instead dependent on post-synaptic AMPA receptor (α-amino-3-hydroxy-5-methyl-4-isoxazole propionic acid-receptor, AMPAR) responses. AMPAR-mediated mEPSC responses amplitude was also significantly decreased in *Itm2b*^*D*^ rats (Figure 5A, D, E, G). Also in this case, the reduction is gene-dosage dependent *(Itm2b*^*w/w*^ vs. *Itm2b*^*D/w*^, P=0.0016; *Itm2b*^*w/w*^ vs. *Itm2b*^*D/D*^, P=0.0005). Decay time of mEPSC was not significantly affected in *Itm2b*^*D*^ rats compared to littermate controls (Figure 5A, F, G).

Since mEPSCs AMPAR-mediated responses are reduced in amplitude, we measured the AMPA/NMDA ratio in evoked responses. Consistent with the hypothesis that the Danish *Itm2b* mutation impairs AMPAR-mediated responses, the AMPA/NMDA ratio was reduced in Danish KI rats (Figure 5H). This difference was statistically different only between *Itm2b*^*w/w*^ and *Itm2b*^*D/D*^ rats, with *Itm2b*^*D/w*^ rats showing an intermedia phenotype.

Finally, we examined the effect of the pathogenic Danish mutation on paired-pulse facilitation (PPF). PPF is a form of short-term synaptic plasticity that is in part determined by changes in P*r* of glutamatergic synaptic vesicles (48)(41). Facilitation at both 50ms and 200ms interstimulus interval (ISI), was significantly increased in *Itm2b*^*D/D*^ (Figure 5I). Even in this case the changes were gene-dosage-dependent (50ms ISI: *Itm2b*^*w/w*^ vs. *Itm2b*^*D/D*^, P<0.0001; *Itm2b*^*D/w*^ vs. *Itm2bP*^*D/D*^, P=0.0031; 200ms ISI: *Itm2b*^*w/w*^ vs. *Itm2b*^*D/D*^, P=0.0205). Interestingly, also an increase in PPF is consistent with a decrease in P*r*, just like a decrease in mEPSC frequency. Overall, our data indicate that the pathogenic Danish *Itm2b* mutation alters glutamatergic synaptic transmission at excitatory hippocampal SC-CA3>CA1 synapses in peri-adolescent KI rats. These alterations are like those seen in *Itm2b* KO and *Itm2b*^*D*^/*Itm2b*^*B*^ KI adult mice.

## DISCUSSION

The choice of the genetic approach and the model organisms used to model human diseases have major implications on the phenotypic expression of disease-associated genetic mutations. For the last 13 years, our laboratory has modeled AD and AD-like neurodegenerative disorders in mice, using a KI approach (3,49–52). The KI approach was preferred because it generates models genetically faithful to human diseases and make no preconceived assumption about pathogenic mechanisms (except the unbiased genetic one). We have recently extended our KI modeling of familial and sporadic forms of AD and AD-related disorders to rats (38,42,44,53,54), because the rat is better suited for behavioral tests and other procedures that are important when studying neurodegenerative diseases. In addition, gene-expression differences suggest that rats may be advantageous model of neurodegenerative diseases over mice. Alternative spicing of *Mapt* (12–15), which forms NFTs and is mutated in Frontotemporal Dementia (16–23), leads to expression of tau isoforms with three or four microtubule binding domains (3R and 4R, respectively). Adult human and rat brains express both 3R and 4R tau isoforms (24): in contrast, adult mouse brains express only 4R tau (25), suggesting that the rat may be a better model organism for dementias with tauopathy.

To explore early dysfunctions that may underlie initial mechanisms leading to dementia, we studied young KI rats carrying the *Itm2b*^*D*^ Familial Danish dementia mutation. Consistent with the findings in *Itm2b*^*D/D*^ mouse KIs (41), we found that Bri2-ADan maturation is altered and accumulates in *Itm2b*^*D/D*^ primary neurons (Figure 2). Analysis of APP metabolism in peri-adolescent *Itm2b*^*D*^ Ki rats (Figure 3 and 4) only showed subtle but significant changes in Aβ42 and Aβ43 steady-state levels, which were slightly increased and decreased, respectively (Figure 3).

We have previously shown that the Danish and British *ITM2b* mutations lead to reduced glutamatergic neurotransmitter release and AMPAR-mediated responses in adult *Itm2b*^*B*^ and *Itm2b*^*D*^ mice (41). These reductions are like those seen in adult *Itm2b* knock-out mice (41,47). Interestingly, we detected identical, gene dosage-dependent, pre-and post-synaptic glutamatergic transmission changes in the SC pathway of peri-adolescent *Itm2b*^*D*^ rats (Figure 5). More specifically, the frequency of mEPSC and the PPF are significantly decreased and increased, respectively, in *Itm2b*^*D*^ rats, suggesting a pre-synaptic reduction of the Pr of glutamatergic synaptic vesicles. In addition, mEPSCs amplitude and the AMPA/NMDA ratio were both decreased in *Itm2b*^*D*^ rats, suggesting a post-synaptic reduction of AMPAR-mediated responses. Collectively, these data together with our previously published observations, indicate that the synaptic transmission alteration caused by Danish mutation occur early in life, and are neither species nor gene-editing technology-specific. These studies underlie the potential relevance of our studies to functional changes caused by the pathogenic *ITM2b* mutations in humans.

Given the functional and pathological interaction between APP and BRI2 (26–36), it is possible that the presence of human Aβ in the rat model may lead to an earlier manifestation of synaptic plasticity deficit in rat as compared to mice, which express rodent Aβ. Moreover, the evidence that both APP and BRI2 tune excitatory synaptic transmission, and that these functions are altered by pathogenic mutations in both APP and BRI2 (37,41,47,55–57) suggest that early alterations in glutamatergic transmission may underlie initial pathogenic mechanisms in dementia. Future studies will be needed to test these hypotheses.

## EXPERIMENTAL PROCEDURES

### Rats and ethics statement

Rats were handled according to the NIH Ethical Guidelines for Treatment of Laboratory Animals. The procedures were described and approved by the Institutional Animal Care and Use Committee (IACUC) at Rutgers (IACUC, protocol number PROTO201702513).

### Generation of rats expressing the Danish Itm2b mutation (Itm2^*D*^ rats)

The rat *Itm2b* gene (GenBank accession number: NM_001006963.1; Ensembl: ENSRNOG00000016271) is located on rat chromosome 15. It comprises 6 exons, with ATP start codon in exon 1 and TGA stop codon in exon 6. The FDD mutation (TTTAATTTGTTCTTGAACAGTCAAGAAAAACATTAT) KI site in oligo donor was introduced into exon 6, which is the target site by homology-directed repair. A silent mutation (GTG to GTC) was also introduced to prevent the binding and re-cutting of the sequence by Cas9 after homology-directed repair. The detailed procedures are reported in the Supporting Information file.

### Standard RNA-Seq analysis

Total brain RNA from 21 days old *Itm2b*^*D/D*^ and *Itm2b*^*ww/*^ rats (2 male and 2 females per each genotype) was extracted with RNeasy RNA Isolation kit (Qiagen). Standard RNA-Seq procedures and data analysis was performed by Genewiz following proprietary methods (https://cdn2.hubspot.net/hubfs/3478602/NGS/RNA-Seq/GENEWIZ_RNA-Seq_Technical_Specifications_US.pdf). Student’s t-test was used for all analyses, with data presented as mean ± SD.

### Rats brain proteins preparation, Western blots and ELISA

These procedures were performed as previously described (42,54). Briefly, rats were anesthetized with isoflurane and perfused via intracardiac catheterization with ice-cold PBS. Brains were extracted and homogenized with a glass-teflon homogenizer in 250 mM Sucrose, 20 mM Tris-base pH 7.4, 1 mM EDTA, 1mM EGTA plus protease and phosphatase inhibitors (ThermoScientific). All steps were carried out on ice. Homogenates were solubilized with 1% NP-40 for 30 min rotating and spun at 20,000 g for 10 minutes. Supernatants were collected and protein content was quantified by Bradford.

For Western blot analyses, proteins were diluted with PBS and LDS Sample buffer-10% β-mercaptoethanol (Invitrogen NP0007) and 4.5M urea to 1*μ*g/*μ*l, loaded on a 4-12% Bis-Tris polyacrylamide gel (Biorad 3450125), and transferred onto nitrocellulose at 25V for 7min using the Trans-blot Turbo system (Biorad). Blotting efficiency was visualized by red Ponceau staining on membranes. For Dot-blot analysis 2.5 *μ*g of material was directly spotted with a p20 pipette on a nitrocellulose membrane. Dot membrane was also visualized by red Ponceau after it was totally dried. Membranes were blocked in 5%-milk (Biorad 1706404) for 30 minutes and washed in PBS/Tween20-0.05%. Primary antibodies were applied dilution in blocking solution (Thermo 37573). The following antibodies were used: Polyclonal anti-Bri2 serum test bleeds provided by Cell Signaling Technology was used at 1:500 with overnight shaking at 4°C. APP-Y188 (Abcam32136), Oligomer Aβ A11 (shared by Rakez Kayed’s Lab), LC3A (CST 4599) and LC3B (CST 2775) were used at 1:1000 with same other condition. Secondary antibodies [either anti-mouse (Southern Biotech 1031-05) or a 1:1 mix of anti-rabbit (Southern Biotech, OB405005) and antirabbit (Cell Signaling, 7074)], were diluted 1:1000 in 5% milk and used against either mouse or rabbit primary antibodies for 1 hour at RT, with shaking. Membranes were washed with PBS/Tween20-0.05% (3 times, 10 minutes each time), developed with West Dura ECL reagent (Thermo, PI34076) and visualized on a ChemiDoc MP Imaging System (Biorad). Signal intensity was quantified with Image LAβ software (Biorad). Data were analyzed using Prism software and represented as mean ± SD.

For analysis of human Aβ peptides and sAPPα/sAPPβ, brain lysates were diluted at 4μg/μl. Aβ38, Aβ40, and Aβ42 were measured with V-PLEX Plus Aβ Peptide Panel 1 6E10 (K15200G, Meso Scale Discovery) and sAPPα/sAPPβ were measured with sAPPα/sAPPβ (K15120E, Meso Scale Discovery). Plates were read on a MESO QuickPlex SQ 120. Aβ43 was quantified using the IBL human Aβ43 Assay Kit #27710.

### Primary hippocampal neuron culture

Rat hippocampal neurons were prepared from *Itm2b*^*wlw*^ and *Itm2b^*D,D*^* post-natal day 1 pups. Briefly, after removal of meninges, the hippocampi were collected in HBSS without magnesium and calcium, 1mM sodium pyruvate, 0.1%glucose, 10mM HEPES. Hippocampi were dissected into single cell by trituration followed by 15 minutes incubation at 37°C in 0.25% trypsin. Cells were subsequently treated with 0.1% DNAse (Sigma, dn25) in plating media (BME, 10% FBS, 0.09% Glucose, 1mM Sodium Pyruvate, 2mM Glutamine, 1x Pen/Strep). Cells were filtered through a Falcon 70μm nylon cell strainer and were plated in Poly L lysine pretreated 12-well-plate (300,000 cells/well) in Neurobasal media, 1x B-27, 2mM glutamine, 1x Pen/Strep. Half of the culture media was changed every 2 days.

### Pharmacological treatment and sample preparation

After 9 days in culture, primary neurons were treated with 50μM Chloroquine (Cell signaling, 14774s) or PBS (Veh) for 18 hours. After treatment, cells were washed with PBS and lysed in RIPA buffer with protease/phosphatase inhibitor for 15 minutes on ice. Lysed cells were centrifuged at full speed for 15 minutes. Cell lysates were quantified and analyzed by Western blot as described earlier for brain lysates.

### Electrophysiological recording

These procedures were performed as previously described (44). Briefly, rats were anesthetized with isoflurane and perfused intracardially with an ice-cold cutting solution containing (in mM) 120 choline chloride, 2.6 KCl, 26 NaH CO_3_, 1.25 NaH_2_PO_4_, 0.5 CaCl_2_, 7 MgCl_2_, 1.3 ascorbic acid, 15 glucose, prebubbled with 95% O_2_/5% CO_2_ for 15 min. The brains were rapidly removed from the skull and coronal brain slices containing the hippocampal formation (350μm thick) were prepared in the ice-cold cutting solution bubbled with 95%O_2_/5% CO_2_ using Vibratome VT1200S (Leica Microsystems) and then incubated in an interface chamber in ACSF containing (in mM): 126 NaCl, 3 KCl, 1.2 NaH_2_P0_4_; 1.3 MgCl_2_, 2.4 CaCl_2_, 26 NaHCO_3_, and 10 glucose (at pH 7.3), bubbled with 95% O_2_ and 5% CO_2_ at 30°C for 1hr and then kept at room temperature. The hemi-slices were transferred to a recording chamber perfused with ACSF at a flow rate of ~2ml/min using a peristaltic pump. Experiments were performed at 28.0 ± 1°C.

Whole-cell recordings in the voltage-clamp mode(−70 mv) were made with patch pipettes containing (in mM): 132.5 Cs-gluconate, 17.5 CsCl, 2 MgCl_2_, 0.5 EGTA, 10 HEPES, 4 ATP, and 5 QX-314, with pH adjusted to 7.3 by CsOH. Patch pipettes (resistance, 8-10 MΩ) were pulled from 1.5 mm thin-walled borosilicate glass (Sutter Instruments, Novato, CA) on a horizontal puller (model P-97; Sutter Instruments, Novato, CA). Basal synaptic responses were evoked at 0.05 Hz by electrical stimulation of the Schaffer collateral afferents using concentric bipolar electrodes. CA1 neurons were viewed under upright microscopy (FN-1, Nikon Instruments, Melville, NY) and recorded with Axopatch-700B amplifier (Molecular Devices, San Jose, CA). Data were low-pass filtered at 2 kHz and acquired at 5–10 kHz. The series resistance (Rs) was consistently monitored during recording in case of reseal of ruptured membrane. Cells with Rs >20 MΩ or with Rs deviated by >20% from initial values were excluded from analysis. Excitatory postsynaptic currents (EPSCs) were recorded in ACSF containing the GABA-A receptors inhibitor bicuculline methiodide (15 μM). The stimulation intensity was adjusted to evoke EPSCs that were 40% of the maximal evoked amplitudes (“test intensity”). 5-10 min after membrane rupture, EPSCs were recorded for 7 minutes at a test stimulation intensity that produced currents of ~40% maximum. For recording of paired-pulse ratio (PPR), paired-pulse stimuli with 50ms or 200ms inter-pulse interval were given. The PPR was calculated as the ratio of the second EPSC amplitude to the first. For recording of AMPA/NMDA ratio, the membrane potential was firstly held at−70 mV to record only AMPAR current for 20 sweeps with 20s intervals. Then the membrane potential was turned to +40 mV to record NMDAR current for 20 sweeps with perfusion of 5μM NBQX to block AMPAR. Mini EPSCs were recorded by maintaining neurons at −70 mV with ACSF containing action potentials blocker (1μM TTX) and GABA-A receptors inhibitors (15μM bicuculline methiodide). mEPSCs were recorded for ~10 mins. Data were collected with Axopatch 700B amplifiers and analyzed with pCLAMP10 software (Molecular Devices). mEPSCs are analyzed using mini-Analysis Program.

### Statistics

All the experiments mentioned in the paper were analyzed by one-way ANOVA or two-way ANOVA as indicated. Data showing statistical significance by oneway ANOVA or two-way ANOVA were subsequently analyzed by either Tukey’s multiple comparisons test or Sidak’s multiple comparisons. All statistical analyses were performed using Prism 9 (GraphPad) software.

## Acknowledgement

All authors read and approved the final manuscript.

## Declarations

The datasets used and/or analyzed during the current study are available from the corresponding author on reasonable request.

## Author Contributions

LD generated the animals; KAN set up breeding and genotyped animals; TY performed the biochemical and molecular experiments; WY performed the electrophysiology experiments; All authors designed the experiments; LD and TY wrote the paper.

*L.D. was funded by the NIH/NIA R01AG063407, RF1AG064821, 1R01AG033007*

## References

1. Tamayev, R., Matsuda, S., Fa, M., Arancio, O., and D’Adamio, L. (2010) Danish dementia mice suggest that loss of function and not the amyloid cascade causes synaptic plasticity and memory deficits. Proc Natl Acad Sci U S A 107, 20822–20827.

2. Tamayev, R., Giliberto, L., Li, W., d’Abramo, C., Arancio, O., Vidal, R., and D’Adamio, L. (2010) Memory deficits due to familial British dementia BRI2 mutation are caused by loss of BRI2 function rather than amyloidosis. J Neurosci 30, 14915–14924.

3. Giliberto, L., Matsuda, S., Vidal, R., and D’Adamio, L. (2009) Generation and initial characterization of FDD knock in mice. PLoS One 4, e7900

4. Deacon, R. M. (2006) Housing, husbandry and handling of rodents for behavioral experiments. Nat Protoc 1, 936–946.

5. Whishaw, I. Q., Metz, G. A., Kolb, B., and Pellis, S. M. (2001) Accelerated nervous system development contributes to behavioral efficiency in the laboratory mouse: a behavioral review and theoretical proposal. Dev Psychobiol 39, 151–170.

6. Kepecs, A., Uchida, N., Zariwala, H. A., and Mainen, Z. F. (2008) Neural correlates, computation and behavioural impact of decision confidence. Nature 455, 227–231.

7. Foote, A. L., and Crystal, J. D. (2007) Metacognition in the rat. Curr Biol 17, 551–555.

8. Bartelle, B. B., Barandov, A., and Jasanoff, A. (2016) Molecular fMRI. J Neurosci 36, 4139–4148.

9. Zimmer, E. R., Leuzy, A., Bhat, V., Gauthier, S., and Rosa-Neto, P. (2014) In vivo tracking of tau pathology using positron emission tomography (PET) molecular imaging in small animals. Transl Neurodegener 3, 6

10. Leuzy, A., Zimmer, E. R., Heurling, K., Rosa-Neto, P., and Gauthier, S. (2014) Use of amyloid PET across the spectrum of Alzheimer’s disease: clinical utility and associated ethical issues. Amyloid 21, 143–148.

11. Zimmer, E. R., Leuzy, A., Gauthier, S., and Rosa-Neto, P. (2014) Developments in Tau PET Imaging. Can J Neurol Sci 41, 547–553.

12. Andreadis, A. (2005) Tau gene alternative splicing: expression patterns, regulation and modulation of function in normal brain and neurodegenerative diseases. Biochim Biophys Acta 1739, 91–103.

13. Janke, C., Beck, M., Stahl, T., Holzer, M., Brauer, K., Bigl, V., and Arendt, T. (1999) Phylogenetic diversity of the expression of the microtubule-associated protein tau: implications for neurodegenerative disorders. Brain Res Mol Brain Res 68, 119–128

14. Hong, M., Zhukareva, V., Vogelsberg-Ragaglia, V., Wszolek, Z., Reed, L., Miller, B. I., Geschwind, D. H., Bird, T. D., McKeel, D., Goate, A., Morris, J. C., Wilhelmsen, K. C., Schellenberg, G. D., Trojanowski, J. Q., and Lee, V. M. (1998) Mutation-specific functional impairments in distinct tau isoforms of hereditary FTDP-17. Science 282, 1914–1917.

15. Roberson, E. D., Scearce-Levie, K., Palop, J. J., Yan, F., Cheng, I. H., Wu, T., Gerstein, H., Yu, G. Q., and Mucke, L. (2007) Reducing endogenous tau ameliorates amyloid beta-induced deficits in an Alzheimer’s disease mouse model. Science 316, 750–754.

16. Spillantini, M. G., and Goedert, M. (1998) Tau protein pathology in neurodegenerative diseases. Trends Neurosci 21, 428–433.

17. Goedert, M., Crowther, R. A., and Spillantini, M. G. (1998) Tau mutations cause frontotemporal dementias. Neuron 21, 955–958.

18. Grundke-Iqbal, I., Iqbal, K., Tung, Y. C., Quinlan, M., Wisniewski, H. M., and Binder, L. I. (1986) Abnormal phosphorylation of the microtubule-associated protein tau (tau) in Alzheimer cytoskeletal pathology. Proc Natl Acad Sci U S A 83, 4913–4917.

19. Hutton, M., Lendon, C. L., Rizzu, P., Baker, M., Froelich, S., Houlden, H., Pickering-Brown, S., Chakraverty, S., Isaacs, A., Grover, A., Hackett, J., Adamson, J., Lincoln, S., Dickson, D., Davies, P., Petersen, R. C., Stevens, M., de Graaff, E., Wauters, E., van Baren, J., Hillebrand, M., Joosse, M., Kwon, J. M., Nowotny, P., Che, L. K., Norton, J., Morris, J. C., Reed, L. A., Trojanowski, J., Basun, H., Lannfelt, L., Neystat, M., Fahn, S., Dark, F., Tannenberg, T., Dodd, P. R., Hayward, N., Kwok, J. B., Schofield, P. R., Andreadis, A., Snowden, J., Craufurd, D., Neary, D., Owen, F., Oostra, B. A., Hardy, J., Goate, A., van Swieten, J., Mann, D., Lynch, T., and Heutink, P. (1998) Association of missense and 5’-splice-site mutations in tau with the inherited dementia FTDP-17. Nature 393, 702–705.

20. Stanford, P. M., Shepherd, C. E., Halliday, G. M., Brooks, W. S., Schofield, P. W., Brodaty, H., Martins, R. N., Kwok, J. B., and Schofield, P. R. (2003) Mutations in the tau gene that cause an increase in three repeat tau and frontotemporal dementia. Brain 126, 814–826.

21. Yasuda, M., Takamatsu, J., D’Souza, I., Crowther, R. A., Kawamata, T., Hasegawa, M., Hasegawa, H., Spillantini, M. G., Tanimukai, S., Poorkaj, P., Varani, L., Varani, G., Iwatsubo, T., Goedert, M., Schellenberg, D. G., and Tanaka, C. (2000) A novel mutation at position +12 in the intron following exon 10 of the tau gene in familial frontotemporal dementia (FTD-Kumamoto). Ann Neurol 47, 422–429.

22. Kowalska, A., Hasegawa, M., Miyamoto, K., Akiguchi, I., Ikemoto, A., Takahashi, K., Araki, W., and Tabira, T. (2002) A novel mutation at position +11 in the intron following exon 10 of the tau gene in FTDP-17. J Appl Genet 43, 535–543.

23. Grover, A., England, E., Baker, M., Sahara, N., Adamson, J., Granger, B., Houlden, H., Passant, U., Yen, S. H., DeTure, M., and Hutton, M. (2003) A novel tau mutation in exon 9 (1260V) causes a four-repeat tauopathy. Exp Neurol 184, 131–140.

24. Hanes, J., Zilka, N., Bartkova, M., Caletkova, M., Dobrota, D., and Novak, M. (2009) Rat tau proteome consists of six tau isoforms: implication for animal models of human tauopathies. J Neurochem 108, 1167–1176.

25. McMillan, P., Korvatska, E., Poorkaj, P., Evstafjeva, Z., Robinson, L., Greenup, L., Leverenz, J., Schellenberg, G. D., and D’Souza, I. (2008) Tau isoform regulation is region- and cell-specific in mouse brain. J Comp Neurol 511, 788–803.

26. Matsuda, S., Giliberto, L., Matsuda, Y., Davies, P., McGowan, E., Pickford, F., Ghiso, J., Frangione, B., and D’Adamio, L. (2005) The familial dementia BRI2 gene binds the Alzheimer gene amyloid-beta precursor protein and inhibits amyloid-beta production. J Biol Chem 280, 28912–28916.

27. Fotinopoulou, A., Tsachaki, M., Vlavaki, M., Poulopoulos, A., Rostagno, A., Frangione, B., Ghiso, J., and Efthimiopoulos, S. (2005) BRI2 interacts with amyloid precursor protein (APP) and regulates amyloid beta (Abeta) production. J Biol Chem 280, 30768–30772.

28. Matsuda, S., Giliberto, L., Matsuda, Y., McGowan, E. M., and D’Adamio, L. (2008) BRI2 inhibits amyloid beta-peptide precursor protein processing by interfering with the docking of secretases to the substrate. J Neurosci 28, 8668–8676

29. Matsuda, S., Matsuda, Y., Snapp, E. L., and D’Adamio, L. (2011) Maturation of BRI2 generates a specific inhibitor that reduces APP processing at the plasma membrane and in endocytic vesicles. Neurobiol Aging 32, 1400–1408

30. Matsuda, S., Tamayev, R., and D’Adamio, L. (2011) Increased A3PP processing in familial Danish dementia patients. J Alzheimers Dis 27, 385–391.

31. Tamayev, R., Matsuda, S., Giliberto, L., Arancio, O., and D’Adamio, L. (2011) APP heterozygosity averts memory deficit in knockin mice expressing the Danish dementia BRI2 mutant. EMBO J 30, 2501–2509.

32. Tamayev, R., and D’Adamio, L. (2012) Inhibition of y-secretase worsens memory deficits in a genetically congruous mouse model of Danish dementia. Mol Neurodegener 7, 19

33. Tamayev, R., and D’Adamio, L. (2012) Memory deficits of British dementia knock-in mice are prevented by Aβ-precursor protein haploinsufficiency. J Neurosci 32, 5481–5485.

34. Tamayev, R., Matsuda, S., Arancio, O., and D’Adamio, L. (2012) β-but not y-secretase proteolysis of APP causes synaptic and memory deficits in a mouse model of dementia. EMBO Mol Med 4, 171–179

35. Lombino, F., Biundo, F., Tamayev, R., Arancio, O., and D’Adamio, L. (2013) An intracellular threonine of amyloid-β precursor protein mediates synaptic plasticity deficits and memory loss. PLoS One 8, e57120

36. Biundo, F., Ishiwari, K., Del Prete, D., and D’Adamio, L. (2016) Deletion of the γ-secretase subunits Aph1B/C impairs memory and worsens the deficits of knock-in mice modeling the Alzheimer-like familial Danish dementia. Oncotarget 7, 11923–11944.

37. Tambini, M. D., Yao, W., and D’Adamio, L. (2019) Facilitation of glutamate, but not GABA, release in Familial Alzheimer’s APP mutant Knock-in rats with increased β-cleavage of APP. Aging Cell 18, e13033

38. Tambini, M. D., Norris, K. A., and D’Adamio, L. (2020) Opposite changes in APP processing and human Aβ levels in rats carrying either a protective or a pathogenic APP mutation. Elife 9

39. Vidal, R., Revesz, T., Rostagno, A., Kim, E., Holton, J. L., Bek, T., Bojsen-Møller, M., Braendgaard, H., Plant, G., Ghiso, J., and Frangione, B. (2000) A decamer duplication in the 3’ region of the BRI gene originates an amyloid peptide that is associated with dementia in a Danish kindred. Proc Natl Acad Sci U S A 97, 4920–4925.

40. Choi, S. I., Vidal, R., Frangione, B., and Levy, E. (2004) Axonal transport of British and Danish amyloid peptides via secretory vesicles. FASEB J 18, 373–375.

41. Yin, T., Yao, W., Lemenze, A. D., and D’Adamio, L. (2020) Danish and British dementia ITM2b/BRI2 mutations reduce BRI2 protein stability and impair glutamatergic synaptic transmission. J Biol Chem

42. Tambini, M. D., and D’Adamio, L. (2020) Trem2 Splicing and Expression are Preserved in a Human Abeta-producing, Rat Knock-in Model of Trem2-R47H Alzheimer’s Risk Variant. Sci Rep 10, 4122

43. Ren, S., Breuillaud, L., Yao, W., Yin, T., Norris, K. A., Zehntner, S. P., and D’Adamio, L. (2020) TNF-α-mediated reduction in inhibitory neurotransmission precedes sporadic Alzheimer’s disease pathology in young. J Biol Chem

44. Ren, S., Yao, W., Tambini, M. D., Yin, T., Norris, K. A., and D’Adamio, L. (2020) Microglia *TREM2*^*R47H*^ Alzheimer-linked variant enhances excitatory transmission and reduces LTP via increased TNF-α levels. Elife 9

45. Shankar, G. M., Li, S., Mehta, T. H., Garcia-Munoz, A., Shepardson, N. E., Smith, I., Brett, F. M., Farrell, M. A., Rowan, M. J., Lemere, C. A., Regan, C. M., Walsh, D. M., Sabatini, B. L., and Selkoe, D. J. (2008) Amyloid-beta protein dimers isolated directly from Alzheimer’s brains impair synaptic plasticity and memory. Nat Med 14, 837–842.

46. Kayed, R., Head, E., Thompson, J. L., McIntire, T. M., Milton, S. C., Cotman, C. W., and Glabe, C. G. (2003) Common structure of soluble amyloid oligomers implies common mechanism of pathogenesis. Science 300, 486–489

47. Yao, W., Yin, T., Tambini, M. D., and D’Adamio, L. (2019) The Familial dementia gene ITM2b/BRI2 facilitates glutamate transmission via both presynaptic and postsynaptic mechanisms. Sci Rep 9, 4862

48. Zucker, R. S., and Regehr, W. G. (2002) Short-term synaptic plasticity. Annu Rev Physiol 64, 355–405.

49. Barbagallo, A. P., Wang, Z., Zheng, H., and D’Adamio, L. (2011) The intracellular threonine of amyloid precursor protein that is essential for docking of Pin1 is dispensable for developmental function. PLoS One 6, e18006

50. Barbagallo, A. P., Wang, Z., Zheng, H., and D’Adamio, L. (2011) A single tyrosine residue in the amyloid precursor protein intracellular domain is essential for developmental function. J Biol Chem 286, 8717–8721.

51. Barbagallo, A. P., Weldon, R., Tamayev, R., Zhou, D., Giliberto, L., Foreman, O., and D’Adamio, L. (2010) Tyr(682) in the intracellular domain of APP regulates amyloidogenic APP processing in vivo. PLoS One 5, e15503

52. Garringer, H. J., Murrell, J., D’Adamio, L., Ghetti, B., and Vidal, R. (2010) Modeling familial British and Danish dementia. Brain Struct Funct 214, 235–244

53. Ren, S., Breuillaud, L., Yao, W., Yin, T., Norris, K. A., Zehntner, S. P., and D’Adamio, L. (2020) TNF-α-mediated reduction in inhibitory neurotransmission precedes sporadic Alzheimer’s disease pathology in young Trem2 R^47H^ rats. J Biol Chem 296, 100089

54. Tambini, M. D., and D’Adamio, L. (2020) Knock-in rats with homozygous *PSEN1 L^435F^* Alzheimer mutation are viable and show selective y-secretase activity loss causing low Aβ40/42 and high Aβ43. J Biol Chem 295, 7442–7451.

55. Yao, W., Tambini, M. D., Liu, X., and D’Adamio, L. (2019) Tuning of Glutamate, But Not GABA, Release by an Intrasynaptic Vesicle APP Domain Whose Function Can Be Modulated by β- or α-Secretase Cleavage. J Neurosci 39, 69927005

56. Fanutza, T., Del Prete, D., Ford, M. J., Castillo, P. E., and D’Adamio, L. (2015) APP and APLP2 interact with the synaptic release machinery and facilitate transmitter release at hippocampal synapses. Elife 4, e09743

57. Del Prete, D., Lombino, F., Liu, X., and D’Adamio, L. (2014) APP is cleaved by Bace1 in pre-synaptic vesicles and establishes a presynaptic interactome, via its intracellular domain, with molecular complexes that regulate pre-synaptic vesicles functions. PLoS One 9, e108576

